# CD45-antibody-drug conjugate clears tissue resident myeloid cells from their niches enabling therapeutic adoptive cell transfer

**DOI:** 10.1101/2023.09.05.556397

**Authors:** Karin Gustafsson, Catherine Rhee, Vanessa Frodermann, Elizabeth W. Scadden, Dan Li, Yoshiko Iwamoto, Rahul Palchaudhuri, Sharon L. Hyzy, Anthony E. Boitano, Matthias Nahrendorf, David T. Scadden

## Abstract

Tissue resident myeloid cells (TRM) in adults have highly variable lifespans and may be derived from early embryonic yolk sac, fetal liver or bone marrow. Some of these TRM are known pathogenic participants in congenital and acquired diseases. Myeloablative conditioning and hematopoietic stem cell transplant can replace long-lived brain TRM resulting in clinical improvements in metabolic storage diseases. With the advent of antibody-drug-conjugate (ADC) targeted cell killing as a cell selective means of transplant conditioning, we assessed the impact of anti-CD45-ADC on TRM in multiple tissues. Replacement of TRM ranged from 40 to 95 percent efficiencies in liver, lung, and skin tissues, after a single anti-CD45-ADC dose and bone marrow hematopoietic cell transfer. Of note, the population size of TRM in tissues returned to pre-treatment levels suggesting a regulated control of TRM abundance. As expected, brain, microglia were not affected, but brain monocytes and macrophages were 50% replaced. Anti-CD45-ADC and adoptive cell transfer were then tested in the chronic acquired condition, atherosclerosis exacerbated by *Tet2* mutant clonal hematopoiesis. Plaque resident myeloid cells were efficiently replaced with anti-CD45-ADC and wild-type bone marrow cells. Notably, this reduced existent atherosclerotic plaque burden. Overall, these results indicate that anti-CD45-ADC clears both HSC and TRM niches enabling cell replacement to achieve disease modification in a resident myeloid cell driven disease.

## Introduction

Antibody-drug-conjugates (ADCs) may be an efficient way of depleting endogenous hematopoiesis to make way for an incoming cell graft while circumventing many of the liabilities of broad cytotoxic interventions like irradiation or chemotherapy. ADC mediated conditioning allows for selective delivery of a toxic payload, limiting off target effects and has been shown to minimize the duration of immunodeficiency^1–4^. However, due to the biodistribution properties of antibodies questions have been raised as to its potential application to conditions where mature leukocytes, not just bone marrow residing hematopoietic stem and progenitors (HSPCs), need to be replaced^5^.

Many tissues contain myeloid cell populations that are bone marrow independent, so-called tissue resident myeloid cells (TRM). Brain microglia are descendants of the first wave of hematopoietic progenitors that originate in the yolk sac^6,7^. In other organs such as liver, lung and skin tissues TRM are primarily derived from later fetal hematopoietic waves^8,9^. These tissue specific myeloid cells will mostly remain bone marrow independent throughout life unless tissue homeostasis is disrupted^10^. Liver Kupffer cells for instance can be depleted by inflammatory injury, leading to the subsequent recruitment of bone marrow derived monocytes that will eventually differentiate into Kupffer cells^11,12^. Likewise, while initially monocyte-derived, the majority of macrophages in advanced atherosclerotic plaques self-renew to maintain their numbers rather than relying on continuous import of cells from the bone marrow^13^. Conventional HSCT conditioning regimens replace TRM efficiently in most tissues, including the central nervous system ^10,14–18^. Antibodies, however, will penetrate solid tissues to varying degrees due to their large size and polarity. The sinusoidal vessel network of the bone marrow leads to efficient antibody uptake but other tissue types with more restrictive vascular beds such as skin or brain will see a more limited distribution. An ADC that facilitates excellent bone marrow conditioning and HSPC engraftment could thus still fail to replace some tissue resident myeloid cell populations.

We set out to test whether a newly developed ADC targeting the leukocyte-restricted, pan-hematopoietic marker CD45 would be able to also deplete tissue resident populations of myeloid cells leading to their replacement upon bone marrow cell transfer. We demonstrate that CD45-ADC and bone marrow cell infusion can lead to complete or near complete substitution of myeloid cells in skin, lung, and liver. Furthermore, in a model of clonal hematopoiesis and atherosclerosis, CD45-ADC treatment followed by bone marrow cells leads to highly efficient depletion of disease driving leukocytes in both bone marrow and atherosclerotic plaques and to disease mitigation.

## Results

### CD45-ADC conditioning and tissue chimerism

We first assessed the efficiency of CD45-ADC treatment and HSCT in replacing tissue resident myeloid cells in WT C57Bl/6 mice. The mice were treated with 3 mg/kg CD45-ADC or Isotype-ADC followed by transplantation of 2.0x 10^7^ congenic bone marrow cells from *Ubiquitin-GFP* donor mice (Figure 1A). As described previously, this resulted in depletion of endogenous cells and rapid engraftment of donor hematopoietic cells in blood and bone marrow 6 weeks later in mice that received CD45-ADC treatment (Figure 1B-C, Supplemental figure 1A-B)^4^. Recipients that were conditioned with Isotype-ADC showed minimal GFP chimerism (Figure 1B-C, Supplemental figure 1A-B). Long-term hematopoietic stem cells (LT-HSCs) as well as myeloid progenitors (megakaryocytic-erythroid progenitors; MEP, common myeloid progenitors; CMP and granulocyte-monocyte progenitors; GMP) all showed close to 90% engraftment (Figure 1C) and numbers of hematopoietic stem and progenitor cells (HSPCs) had largely returned to baseline levels (Supplemental figure 1C). Relative to other immune cells, donor chimerism of peripheral blood T cells was lower (∼60%, Figure 2C), likely due to the slow recovery of T cell production post-HSCT in combination with quiescent, peripheral T cells being less sensitive to the ADC payload^4^.

**Figure 1.**
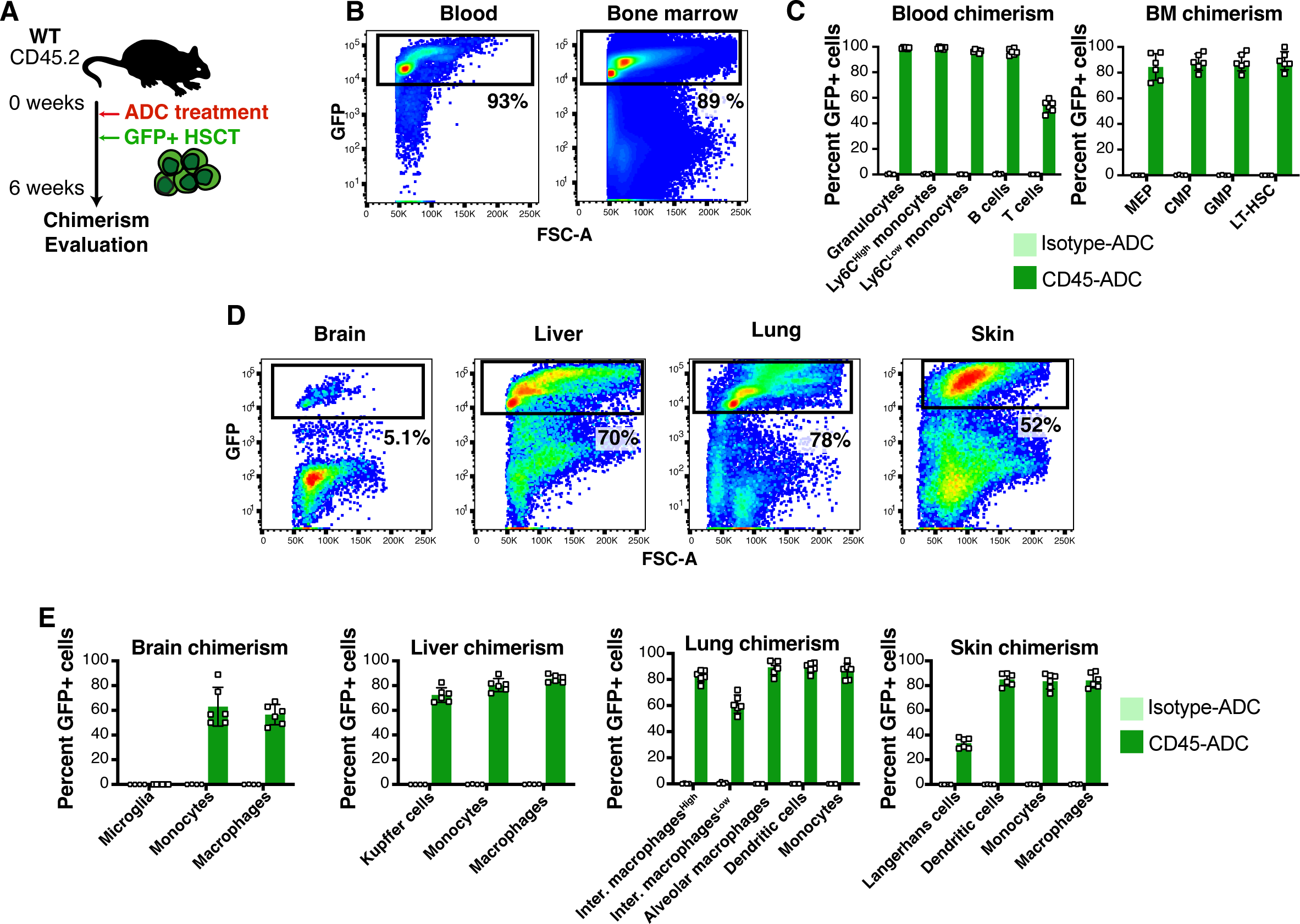
(A) Schematic depiction of the experimental design for CD45-ADC treatment and Ubiquitin-GFP HSCT. (B) Representative flow plots showing the degree of GFP chimerism in peripheral blood and bone marrow 6 weeks post-CD45-ADC treatment and HSCT. (C) Bar graphs displaying the flow cytometric analysis of GFP chimerism in peripheral blood and bone marrow cell subsets 6 weeks following CD45-ADC treatment and HSCT. Megakaryocyte-erythroid progenitor (MEP); Common myeloid progenitor (CMP); Granulocyte-monocyte progenitor (GMP); Long-term hematopoietic stem cell (LT-HSC). (D) Flow cytometric analysis plots representative of hematopoietic GFP chimerism in brain, liver, lung and skin 6 weeks following treatment with CD45-ADC and HSCT. (E) The GFP chimerism in various myeloid cell populations in brain, liver, lung and skin, as assessed by flow cytometry, summarized in bar graphs 6 weeks after CD45-ADC and HSCT. MHCII High interstitial macrophages (Inter. macrophages^High^); (MHCII Low interstitial macrophages (Inter. macrophages^Low^).

**Figure 2.**
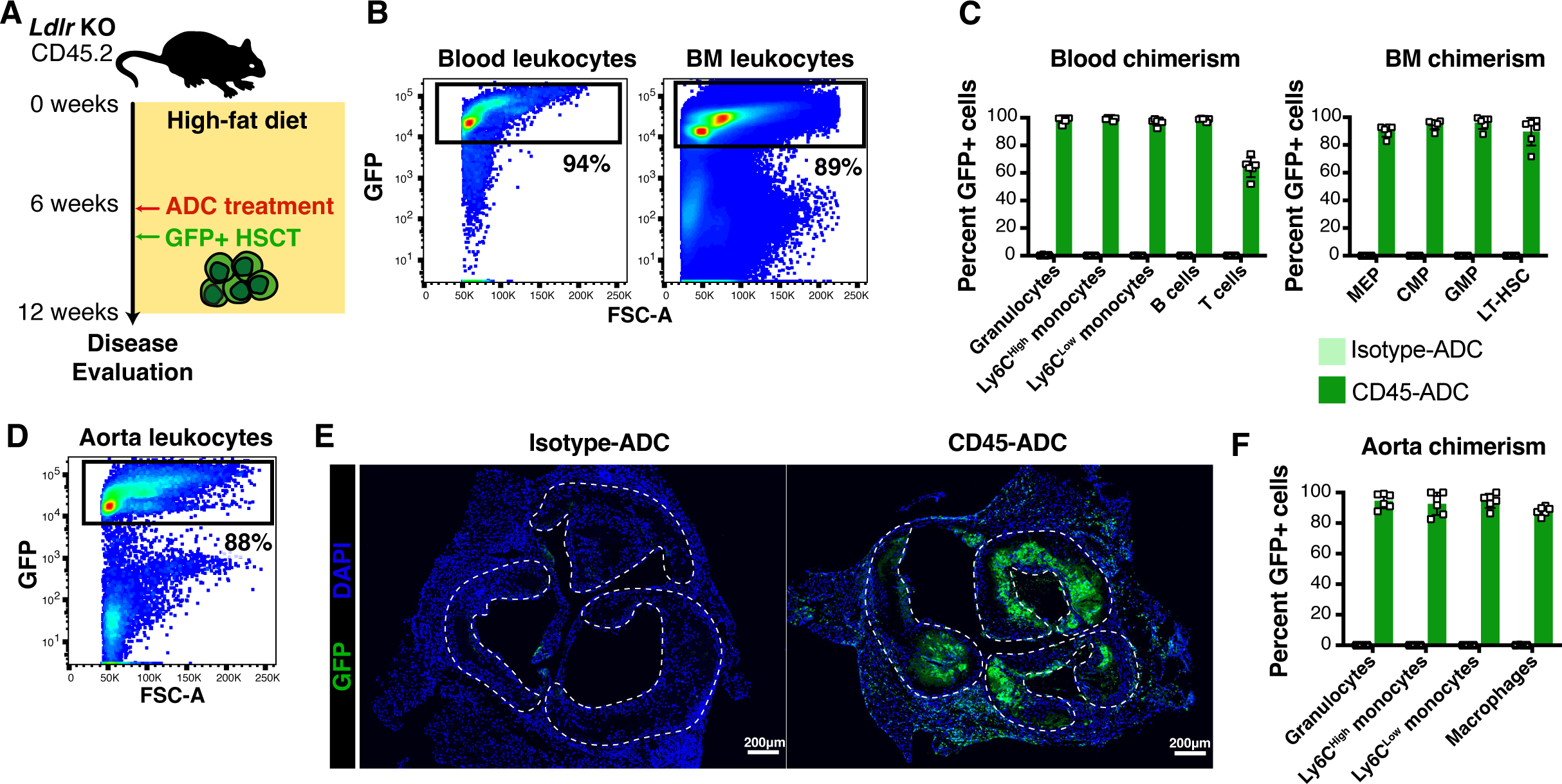
(A) Schematic illustration of the experimental design for atherosclerosis induction and CD45-ADC treatment and Ubiquitin-GFP HSCT in Ldlr KO mice. (B) Representative flow plots showing the degree of GFP chimerism in peripheral blood and bone marrow 6 weeks post-CD45-ADC treatment and HSCT in atherosclerotic Ldlr KO mice. (C) Bar graphs displaying the flow cytometric analysis of GFP chimerism in peripheral blood and bone marrow cell subsets 6 weeks following CD45-ADC treatment and HSCT atherosclerotic Ldlr KO mice. Megakaryocyte-erythroid progenitor (MEP); Common myeloid progenitor (CMP); Granulocyte-monocyte progenitor (GMP); Long-term hematopoietic stem cell (LT-HSC). (D) FACS plot representative of hematopoietic GFP chimerism in aortic atherosclerotic lesions from Ldlr KO mice 6 weeks following treatment with CD45-ADC and HSCT. (E) Immunofluorescence images of aortic roots from Ldlr KO mice stained for GFP (green) and DAPI (blue). Dashed lines outline atherosclerotic plaque surface area. Scale bar 200μm. (F) GFP chimerism in myeloid cell populations infiltrating aortic atherosclerotic plaques 6 weeks after CD45-ADC treatment and HSCT of Ldlr KO presented as bar graphs.

To determine how well various leukocyte types were depleted in non-hematopoietic tissues we also collected brain, liver, lung, and skin from the mice. Flow cytometric analysis of the hematopoietic fraction of these tissues revealed varying degrees of GFP chimerism. While liver and lung displayed more than 70% GFP engraftment after CD45-ADC treatment, average skin GFP chimerism was 52% and as expected brain showed little infiltration of graft derived cells (Figure 1D, Supplemental figures 2A-D and 3A-D). This was further reflected in the analysis of individual myeloid cell types from these tissues. Immunophenotypic microglia that represent the largest population of hematopoietic cells in brain parenchyma (Supplemental figure 2A-B), show little to no GFP labeling following CD45-ADC conditioning (Figure 1E). This is in line with the expectation of minimal uptake of antibodies across the blood-brain barrier^25^. Brain macrophages and monocytes on the other hand are bone marrow derived and show significant GFP engraftment (Figure 1E). For liver, lung, and skin, GFP engraftment was robust in most myeloid lineages following CD45-ADC conditioning, ranging from on average 60-90% (Figure 1E, Supplemental figures 2C-D, and 3A-D). The one exception being the epidermal Langerhans cells that show an average GFP chimerism of 33% (Figure 1E). The epidermis of the skin is avascular, reducing antibody diffusion and thus most likely decreasing Langerhans cell depletion. Notably, bone marrow derived, non-Kupffer cell macrophages are rarely found in the liver at steady state (Supplemental figure 2C-D), while we observed a substantial fraction of non-Kupffer cell macrophages 6 weeks post-CD45-ADC and bone marrow infusion (Supplemental figure 2C-D). Tim4, the marker used for immunophenotypic identification of Kupffer cells, is however one of the last to be upregulated when bone marrow derived cells differentiate into Kupffer cells^26^. Additionally, the combined number of immunophenotypic macrophages and Kupffer cells per liver was not different between CD45- and Isotype-ADC treated mice (Supplemental figure 2E), suggesting that the increased number of macrophages in the CD45-ADC cohort might merely be the result of the bone marrow derived cells not having completed their maturation to Kupffer cells. In summary, this demonstrates that CD45-ADC conditioning and bone marrow hematopoietic cell infusion leads to efficient and rapid replacement of multiple tissue resident myeloid cell populations, many of which are bone marrow independent at homeostasis.

### Near complete replacement of myeloid cells in atherosclerotic plaques following CD45-ADC conditioning and transplantation

Macrophages are drivers of atherosclerotic disease and in advanced atherosclerotic plaques local proliferation, rather than recruitment of bone marrow derived monocytes, leads to their accumulation in atherosclerotic plaques^13^. We therefore tested if HSCT following CD45-ADC conditioning would facilitate replacement of plaque macrophages in a murine model of atherosclerosis. As was done for WT recipient mice, *Ldlr* KO mice with established atherosclerotic lesions were conditioned with 3 mg/kg CD45-ADC or Isotype-ADC and subsequently transplanted with 2.0x 10^7^ congenic *Ubiquitin-GFP* bone marrow cells (Figure 2A). Blood and bone marrow were isolated an additional 6 weeks post-dosing, revealing similar, high levels of GFP engraftment (Figure 2B) and most bone marrow HSPCs and their respective mature lineages displayed more than 90% GFP chimerism in CD45-ADC treated mice compared to Isotype controls (Figure 2C, Supplementary figure 4A). The exception being T lymphocytes that, just as was seen in WT recipients, had notably fewer GFP+ cells (Figure 2C).

Analysis of aortic plaques from these mice demonstrated significant GFP engraftment in CD45-ADC conditioned recipients (Figure 2D and E). Individual myeloid lineages, including plaque macrophages, also showed near complete replacement by GFP+ cells in CD45-ADC conditioned mice (Figure 2F, Supplementary figure 4B). Lastly, the absolute number of myeloid cell types were not different between CD45- and Isotype-ADC treated mice, suggesting that the tissue had returned to homeostasis following the transplantation (Supplementary figure 4C). Conditioning with CD45-ADC followed by bone marrow hematopoietic cell infusion thus enables efficient replacement of aortic plaque macrophages.

### CD45-ADC conditioning and transplantation depletes *Tet2* mutant hematopoietic stem and progenitor cells

Age-related clonal hematopoiesis is associated with atherosclerotic disease^27,28^. Clonal hematopoiesis arises when single LT-HSCs acquire certain mutations, which leads to bone marrow HSPCs producing increased numbers of myeloid progeny and to mature myeloid cells secreting excess amounts of inflammatory cytokines in the case of Tet2 mutation^29^. To model clonal hematopoiesis, we set up bone marrow chimeras in atherosclerosis-prone *Ldlr* KO recipients, where the mice were transplanted with either 20% *Tet2* KO CFP+ (*Tet2^fl/f^MxCre-βactin-CFP)* and 80% WT GFP+ (*Ubiqutin-GFP)* bone marrow or a 20% WT CFP+ (*Tet2^WT/WT^MxCre-βactin-CFP*) and 80% WT GFP+ graft (Figure 3A). Highly efficient *Tet2* deletion was induced at 4 weeks (Supplementary figure 5A) and the mice were subsequently placed on a high-fat diet to initiate atherosclerosis. CD45- or Isotype-ADC (3mg/kg) treatment was given after 6 weeks of disease development, and all mice received 2.0x 10^7^ congenic CD45.1+ bone marrow cells (Figure 3A).

**Figure 3.**
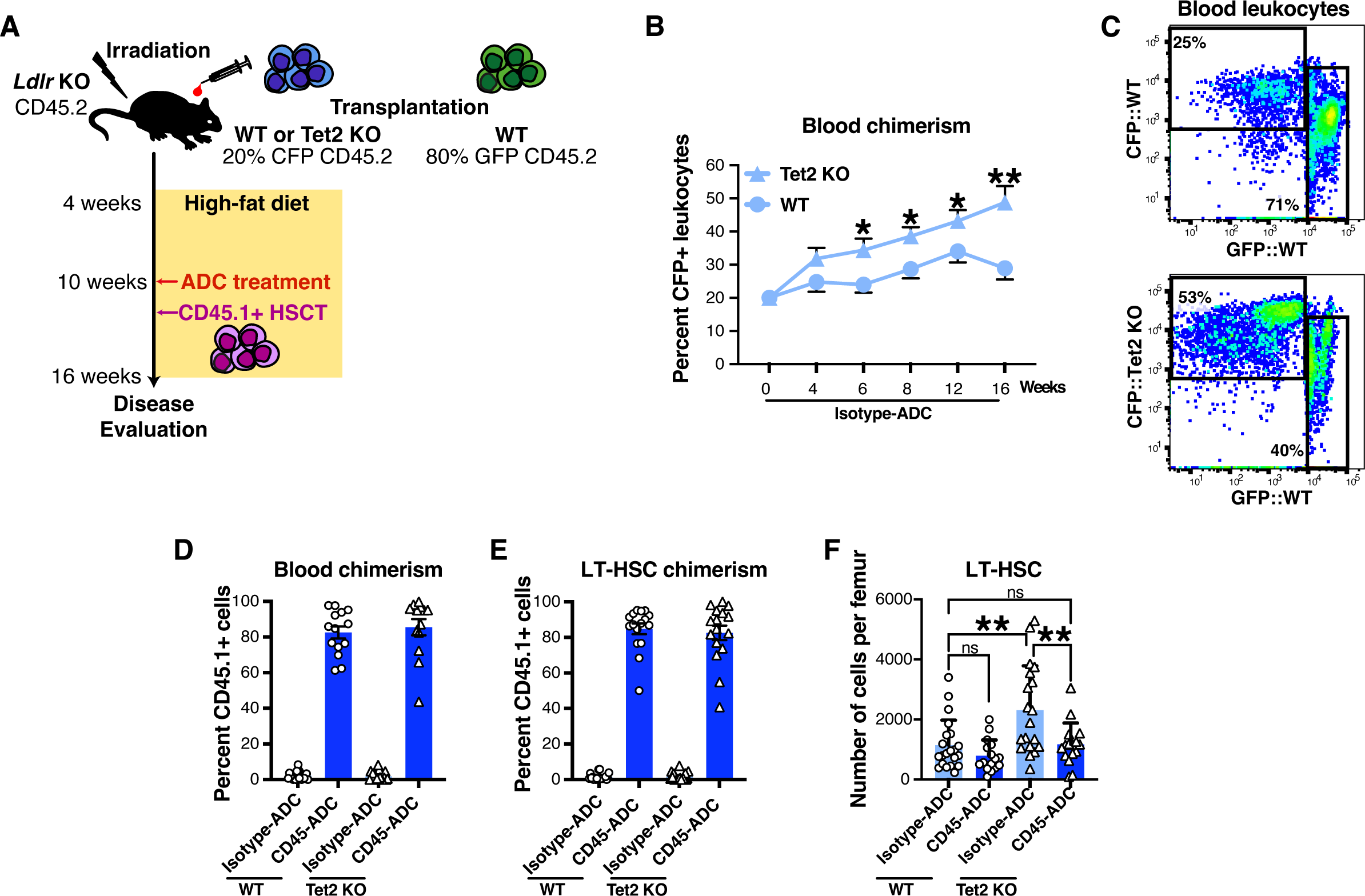
(A) Schematic depiction of the experimental outline for modelling Tet2 KO driven clonal hematopoiesis and atherosclerosis. (B) Peripheral blood WT or Tet2 KO chimerism (CFP) in atherosclerotic Ldlr KO recipient mice weeks 0-16 post-bone marrow transplantation. (C) Representative FACS plots of CFP chimerism in peripheral blood 16 weeks after transplantation into Ldlr KO mice. (D) Bar graphs showing FACS analysis of CD45.1 chimerism 6 weeks after CD45-ADC treatment and CD45.1+ HSCT in atherosclerotic Ldlr KO mice with either WT or Tet2 KO bone marrow. (E) FACS assessment of CD45.1 chimerism 6 weeks after CD45-ADC treatment and CD45.1+ HSCT in atherosclerotic Ldlr KO mice with either WT or Tet2 KO bone marrow displayed as bar graphs. (F) Number of LT-HSCs per femur in atherosclerotic Ldlr KO mice with either WT or Tet2 KO bone marrow as determined by flow cytometric analysis.

*Tet2* KO gives mutant HSPCs a competitive advantage^30^. The proportion of *Tet2* mutant derived cells in peripheral blood consequently increased with time when comparing Isotype control treated *Tet2* KO recipients to their WT counterparts (Figure 3B) and at 16 weeks post-transplantation the average *Tet2* KO chimerism increased more than two-fold (Figure 3B and C). This is also reflected in the bone marrow where the vast majority of LT-HSCs were found to be *Tet2* KO at 16 weeks in the Isotype-ADC cohort (Supplementary figure 5B). Despite this obvious competitive advantage, CD45-ADC and subsequent infusion of a WT bone marrow graft resulted in efficient replacement of *Tet2* KO cells in both blood and LT-HSCs, as evident by the high degree of CD45.1 engraftment (Figure 3C and D). This also led to a normalization of white blood cell counts (Supplemental figure 5C) and numbers of LT-HSCs residing in the bone marrow when comparing *Tet2* KO recipients treated with CD45-ADC to controls (Figure 3E). Most aspects of *Tet2* mutant clonal hematopoiesis in blood and bone marrow were thus reversed by CD45-ADC conditioning and transplantation with WT bone marrow.

### Clonal hematopoiesis associated atherosclerosis is mitigated by CD45-ADC conditioning and WT hematopoietic cells

*Tet2* KO leukocytes also displayed a selective advantage in atherosclerotic plaques isolated from the aorta of WT and *Tet2* mutant recipients treated with Isotype-ADC, 12 weeks post-disease induction (Figure 4A, Supplementary figure 5D). Nonetheless, CD45-ADC and bone marrow hematopoietic cell infusion lead to efficient replacement of *Tet2* KO cells in atherosclerotic lesions 6 weeks following conditioning, as evidenced by the high degree of CD45.1+ chimerism (Figure 4B and C). This also resulted in the normalization of multiple parameters in CD45-ADC treated *Tet2* KO recipients. Overall aorta cellularity was reduced to WT quantities (Supplemental figure 5E) and numbers of Ly6C^High^ monocytes as well as macrophages also returned to levels similar to what was observed in WT controls (Figure 4D and E). Aortic granulocyte and Ly6C^Low^ monocytes remained unchanged across genotype and treatments (Figure 4F and G). We next assessed aortic plaque size as a measure of disease burden. As has been described by others, we observed that *Tet2* mutant leukocytes contributed to increased atherosclerotic plaques in aortic roots compared to WT controls (Figure 5A and B). Plaque burden was however specifically reduced in *Tet2* KO recipients treated with CD45-ADC and WT bone marrow cells (Figure 5A and B), suggesting that clonal hematopoiesis driven atherosclerosis can be reversed if mutant HSPCs and mature myeloid leukocytes are replaced by WT cells.

**Figure 4.**
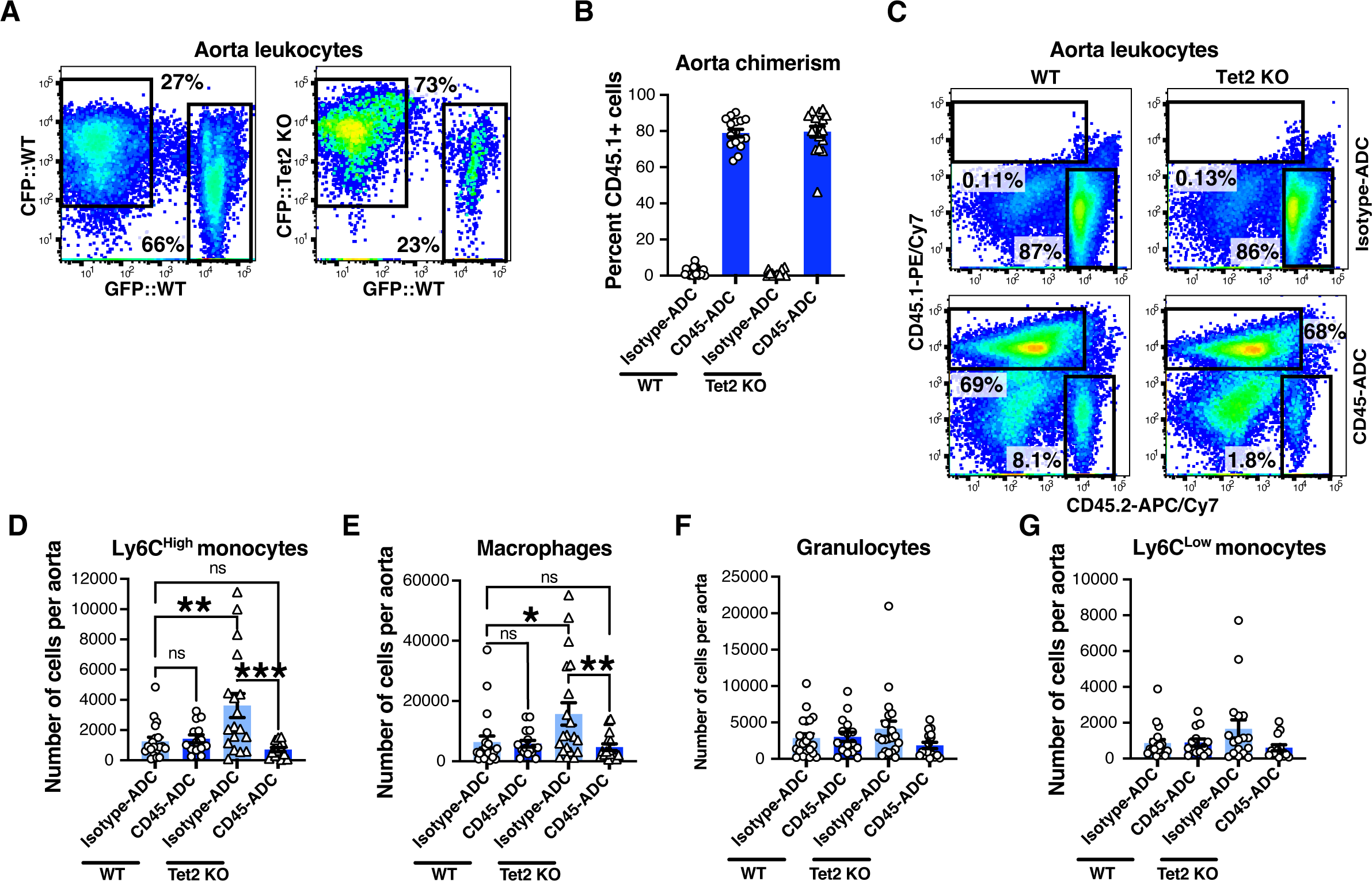
(A) Representative flow plots showing the degree of CFP chimerism in atherosclerotic aortic lesions 16 weeks after transplantation into Ldlr KO mice. (B) Bar graphs showing flow cytometric analysis of CD45.1 chimerism 6 weeks after CD45-ADC treatment and CD45.1+ HSCT in atherosclerotic Ldlr KO mice with either WT or Tet2 KO bone marrow. (C) Flow plots representative of atherosclerotic aortic lesion CD45.1 chimerism post-CD45-ADC treatment and CD45.1+ HSCT in Ldlr KO mice with either WT or Tet2 KO bone marrow. (D-G) Absolute numbers of various myeloid immune cells in aortic atherosclerotic plaques 6 weeks after CD45-ADC treatment and CD45.1+ HSCT in Ldlr KO mice with either WT or Tet2 KO bone marrow as analyzed by flow cytometry.

**Figure 5.**
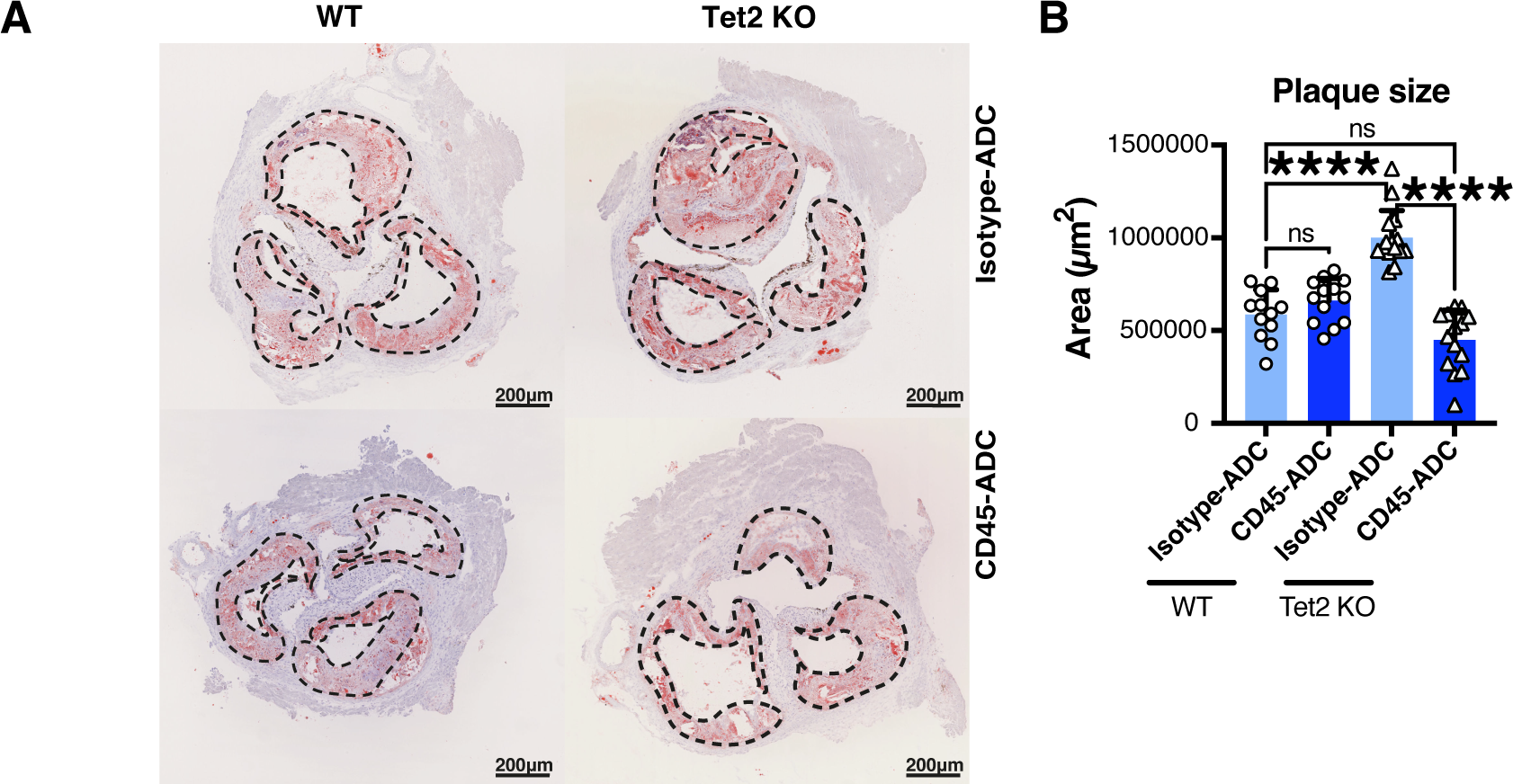
(A) Oil red O images of aortic roots, 6 weeks after CD45-ADC treatment and CD45.1+ HSCT in Ldlr KO mice with either WT or Tet2 KO bone marrow. Dashed lines outline atherosclerotic plaque surface area. Scale bar represents 200μm. (B) Quantification of aortic root atherosclerotic lesion surface area 6 weeks after CD45-ADC treatment and CD45.1+ HSCT in Ldlr KO mice with either WT or Tet2 KO bone marrow

## Discussion

TRM are key participants in tissue homeostasis and as such may also be drivers of disease. A wide range of diseases can thus be cured if TRMs could be replaced efficiently. We report that a single dose of reduced toxicity ADC conditioning and adoptive hematopoietic cells can replace TRM and enable a therapeutic effect..

HSCT is a curative alternative for metabolic storage diseases like Hurler syndrome or Gaucher disease, as it is well established that radiation or chemotherapy conditioning ablates native TRM, allowing for replacement of the disease-causing cells upon transplant ^31,32^. However, due to the morbidity and mortality associated with standard conditioning regimens many patients do not receive this treatment. Instead, they may rely on lifelong, costly and time-consuming enzyme replacement therapies, if they are available. Yet, previous experimental work suggests that only 10% of the mutant macrophages need to be replaced to achieve disease modification^33^. CD45-ADC conditioning followed by bone marrow infusion led to a minimum of 30% TRM engraftment efficiency but for most peripheral tissues TRM replacement was 80-90%. One major limitation is the absence of microglial replacement, though approximately half of central nervous system (CNS) monocytes and macrophages were from the graft. Whether those cells would be sufficient to enable a therapeutic effect is unknown.

Previous work in mouse models using radiation conditioning has shown that host derived TRM will repopulate tissues if the incoming HSCT graft has an inherent deficiency in macrophage maintenance^10^. It was therefore unclear whether WT derived myeloid cells would be able to outcompete *Tet2* mutant macrophages in atherosclerotic lesions given their competitive disadvantage^29^. *Tet2* KO macrophage replacement was however nearly complete and resulted in a significant reduction in atherosclerotic plaque burden, suggesting that CD45-ADC treatment is highly efficient at ablating TRM. There are currently no treatments that target clonal hematopoiesis, but these results suggest that some form of TRM replacement strategy might be helpful.

For both metabolic storage disease and clonal hematopoiesis exacerbated atherosclerosis, complete replacement of the hematopoietic system would be an extreme measure, but it may not be needed. It is reported that most TRM become self-sustaining once established in their respective tissues^13,15^. Follow up studies will determine whether durable TRM chimerism can be achieved with lower doses of ADC and only mature or progenitor level myeloid cell infusions. If so, it is possible to envision using ADC to ablate mutant host-derived cells and follow up with infusion of gene corrected myeloid progenitors or mature macrophages. Indeed, in animal models of hereditary pulmonary alveolar proteinosis disease causing *CSF2RA/B* mutant macrophages can be durably replaced by WT macrophages, leading to disease correction^34^. Our data suggest that the CD45-ADC may enable such an approach for additional conditions.

Remarkably, in all tested tissues where TRM replacement occurred post-CD45-ADC conditioning, cell numbers returned to baseline levels (Supplementary figures 2B, D, and E, 3B and D, 4C). A similar phenomenon was observed in the aortic plaques. When macrophage numbers were normalized to plaque size, there was no difference across genotypes or treatment groups (Supplementary figure 5F). The latter observation is in line with what others have reported in the context of *Tet2* mutant atherosclerosis^29^. This supports the idea that the surrounding tissue regulates the size of its TRM population^35^, perhaps by TRM ‘niches’ regulating TRM persistence in a highly restricted manner.

In summary, this study provides proof-of concept that targeted, reduced toxicity CD45-ADC conditioning and hematopoietic cell adoptive transfer can be used to replace TRM achieving a therapeutic effect in clonal hematopoiesis driven atherosclerotic disease. Furthermore, in other diseases where TRM participate, a similar strategy can be envisioned by using TRM replacement with either WT or perhaps hematopoietic cells expressing therapeutic proteins in other disease settings where TRMs participate.

## Methods

### Antibody-drug-conjugate (ADC)

The CD45-ADC and its cognate Isotype Control-ADC have been reported previously (Magenta Therapeutics)^4^. In brief, CD45.2 clone 104 was engineered for increased *in vivo* clearance and conjugated to the pyrrolobenzodiazepine (PBD) cytotoxic payload.

### Animals

ADC conditioning and hematopoietic stem cell transplantation (HSCT) experiments in wild type (WT) mice were carried out in 8-week-old male and female C57BL/6J mice. For modelling of atherosclerotic disease 8-week-old make and female B6.129S7-Ldlrtm1Her/J *low density lipoprotein receptor* (*Ldlr*) knockout (KO) female and male mice were used. Donors of GFP labeled WT marrow were C57BL/6-Tg(UBC-GFP)30Scha/J mice and B6.SJL-Ptprca Pepcb/BoyJ were used as CD45.1 donors. *Tet2* KO mouse strain B6;129S-Tet2tm1.1Iaai/J was crossed to B6.Cg-Tg(Mx1-cre)1Cgn/J and B6.129(ICR)-Tg(CAG-ECFP)CK6Nagy/J to generate CFP labeled conditional *Tet2* WT and KO donors for clonal hematopoiesis modelling. All mice were obtained from Jackson Laboratories and all animal experimentation was carried out in accordance with national and institutional guidelines and with the approval of the Institutional Animal Care and Use Committee of Massachusetts General Hospital.

### ADC-conditioning and HSCT

Recipient mice, unless otherwise specified, received retro-orbital injections of Isotype-ADC or CD45-ADC (3 mg/kg body weight) and 48-72 hours post-ADC administration 2.0x 10^7^ congenic bone marrow cells from donor mice were injected retro-orbitally. Donor bone marrow was prepared by CO_2_ asphyxiation of *Ubiquitin-GFP* or CD45.1+ donor mice, followed by dissection of femurs, tibias, pelvis, and vertebral column. Bone marrow cells were isolated by crushing the bones gently in phosphate buffered saline (PBS, ThermoFisher Scientific) supplemented with 2% (v/v) fetal bovine serum (FBS, Gibco) and passing the resultant cell suspension over a 70μm cell strainer (BD Falcon). Recipient mice were euthanized for tissue isolation 6 weeks post-HSCT.

### Modelling clonal hematopoiesis

*Ldlr* KO recipient mice received a split dose of 12 Gray irradiation from a ^137^Cs source 24 hours before graft injection. Donor bone marrow was prepared as described above. The graft consisted of either 200 000 *Tet2^fl/f^MxCre-βactin-CFP* and 800 000 *Ubiquitin-GFP* bone marrow cells or 200 000 *Tet2^WT/WT^MxCre-βactin-CFP* 800 000 *Ubiquitin-GFP* bone marrow cells and was delivered via retro-orbital injection. Cre mediated deletion of the floxed *Tet2* allele was induced 4 weeks post-transplantation by 4 injections every other day of 12.5 μg/ml polycytidylic acid (poly:IC, Cytiva).

### Atherosclerosis modelling

To induce atherosclerosis, *Ldlr* KO mice were placed on rodent atherogenic (1.25% cholesterol, regular casein, Research Diets Inc) for a total of 12 weeks. In *Tet2* mutant clonal hematopoiesis experiments, mice were allowed to recover for 4 weeks before being placed in the atherogenic diet.

### Blood sampling

Depending on the experiment, mice were bled at 4, 6, 8, 12 and 16 weeks post-transplant or at 6 weeks post-transplant. All blood collection was performed under isoflurane anesthesia, via retro-orbital puncture and blood was collected into EDTA coated tubes to prevent coagulation. Complete blood counts were obtained using the Element HT5 Auto Hematology system (Heska).

### Tissue collection

For all tissue harvests, the mice were anesthetized using vaporized isoflurane (3-4%) and anesthesia depth was assessed by toe pinching. Transcardial perfusion was thereafter performed with 40 ml PBS (ThermoFisher Scientific). For isolation of lungs 5-10 ml of PBS (ThermoFisher Scientific) was also flushed through the right ventricle. The liver was slowly perfused with an additional 30 ml of PBS (ThermoFisher Scientific), slowly perfused through the portal vein.

Skin was isolated from mouse ears^19^. In brief, ears were excised 2 mm above their base. The internal and external faces of the ear was then separated gently by forceps and placed, cartilage side down in PBS (ThermoFisher Scientific) with 50 mM HEPES (Gibco) and 0.4mg/ml (w/v) Dispase II (Gibco). The tissue was incubated for 2 hours at 37°C. Ear cartilage was thereafter scraped off gently and the remaining skin tissue is finely minced and subjected to 2 rounds of digestion in RPMI 1640 (Gibco) with 50 mM HEPES (Gibco), 2 mg/ml (w/v) Collagenase IV (Millipore Sigma) and 6 U/ml DNase I (ThermoFisher Scientific) at 37°C and 750rpm for 30 minutes. The resulting single cell suspension was passed through a 100μm cell strainer (BD Falcon) and thereafter washed with PBS supplemented with 10% (v/v) FBS (Gibco). Finally, ACK lysis (Quality Biological) for 5 minutes at room temperature was used for removal of erythroid cells.

Brains were carefully dissected, cut into small pieces, and further mechanically dissociated using a p1000 micropipette^20^. The tissue was the enzymatically dissociated using Multi-tissue dissociation kit I (Miltenyi) for 30 minutes at 37°C and 750rpm, with intermittent trituration. The cell suspension was filtered over 100μm cell strainer (BD Falcon) and thereafter washed with PBS (ThermoFisher Scientific) supplemented with 10% (v/v) FBS (Gibco). The samples were further cleaned up by Debris removal solution (Miltenyi) and finally red blood cells were lysed using ACK lysis buffer (Quality Biological).

Liver hematopoietic cells were isolated by careful mincing of hepatic tissue followed by enzymatic dissociation in RPMI with 50 mM HEPES (Gibco), 1 mg/ml Collagenase IV and 6 U/ml DNase I (ThermoFisher Scientific) at 37°C and 750rpm for 30 minutes^21,22^. The liver cell suspension was then passed through a100μm cell strainer (BD Falcon) and thereafter washed with RPMI supplemented with 10% (v/v) FBS (Gibco). Hepatocytes were removed by centrifuging samples for 3 minutes at 50*g*, followed by careful collection of supernatant only. Residual erythrocytes were removed by 5 minutes incubation with ACK lysis buffer at room temperature (Quality Biological).

Lungs were dissected and minced before being enzymatically digested for 30 minutes at 37°C and 750rpm with RPMI 1640 supplemented with 50 mM HEPES (Gibco), 400 μg/ml Liberase TM (Roche) and 6 U/ml DNase I (ThermoFisher Scientific)^23,24^. The digested samples were then filtered across 100μm cell strainer (BD Falcon) and thereafter washed with RPMI supplemented with 10% (v/v) FBS (Gibco). Erythrocytes were lysed with ACK lysis buffer (Quality Biological) and 5 minutes incubation at room temperature.

The entire aorta was harvested from the root to the iliac bifurcation^13^. The aortic root was set aside for histology and the rest of the tissue was cut finely and the incubated in PBS supplemented with 450 U/ml collagenase I (Millipore Sigma), 125 U/ml collagenase XI (Millipore Sigma), 6 U/ml DNase I (ThermoFisher Scientific) and 60 U/ml hyaluronidase (Millipore Sigma) for 1 h at 37 °C while shaking at 750rpm. The resulting cell suspension was then passed over a 100μm cell strainer (BD Falcon) and washed with RPMI 1640 (Gibco) supplemented with 10% (v/v) FBS (Gibco).

Bone marrow was isolated from dissected femurs that were gently crushed in PBS with 2% (v/v) FBS and subsequently filtered over a 70μm cell strainer (BD Falcon). Red cell lysis was done by incubating samples with ACK lysis buffer (Quality Biological) for 5 minutes at room temperature.

### FACS analysis of blood

Red blood cells were lysed with ACK lysis buffer (Quality Biological) for 5 minutes at room temperature. Samples were centrifuged at 800*g* for 5 minutes and resuspended in PBS (ThermoFisher Scientific) with 2% (v/v) FBS (Gibco). Samples were blocked for 10 minutes with murine Fc-block (BD Biosciences) at 4°C. The samples were then stained with the blood lineage antibody panel (Table 1) for 30 minutes at 4°C and then washed with PBS (ThermoFisher Scientific) and 2% (v/v) FBS (Gibco). The samples were resuspended in PBS (ThermoFisher Scientific) with 2% (v/v) FBS (Gibco) and 0.25μg 7-Aminoactinomycin D (7AAD, BD Biosciences) and analyzed on a BD FACS Aria III (BD Biosciences).

### FACS analysis of bone marrow

Bone marrow samples were stained with bone marrow HSPC antibody panels I or II (Table 1) and incubated at 4°C for 60 minutes. Samples were washed and subsequently stained with 1.5 μg/ml Streptavidin-BV650 (Biolegend) for 20 minutes at 4°C. The cells were washed and resuspended in PBS (ThermoFisher Scientific) with 2% (v/v) FBS (Gibco) and 0.25μg 7-Aminoactinomycin D (7AAD, BD Biosciences) and analyzed on a BD FACS Aria III (BD Biosciences).

### FACS analysis of tissue resident macrophages

Brain, skin, lung, and liver single cell suspensions were incubated with murine Fc block for 10 minutes at 4°C. The cells were subsequently stained with their respective, tissue specific antibody panels (Table 1) for 45 minutes at 4°C. The samples were washed with PBS (ThermoFisher Scientific) with 2% (v/v) FBS (Gibco). Lung samples were further incubated with 1.5 μg/ml Streptavidin-BV785 (Biolegend) and then washed again. After the final wash, all samples were resuspended in PBS (ThermoFisher Scientific) with 2% (v/v) FBS (Gibco) and 1 μg/ml 4′,6-diamidino-2-phenylindole (DAPI) and analyzed on a BD FACS Aria III (BD Biosciences).

### FACS analysis of aorta

Single cell suspensions of aortic tissue were blocked with murine Fc block for 10 minutes at 4°C. The samples were then stained with aorta antibody panel (Table 1) for 30 minutes at 4°C, followed by a wash in in PBS (ThermoFisher Scientific) with 2% (v/v) FBS (Gibco). Before analysis, the samples were resuspended in PBS (ThermoFisher Scientific) with 2% (v/v) FBS (Gibco) and 1 μg/ml DAPI and analyzed on a BD FACS Aria III (BD Biosciences).

### Histology

Aortic roots were dissected and embedded in Tissue-Tek O.C.T. (Sakura Finetek), snap frozen with dry ice cooled 2-Methylbutane (Fisher Scientific) and subsequently sectioned into 6μm slices. To ensure lesion size was accurately estimated between treatment groups, sections that captured the maximum lesion area were used. Lesion size was visualized by Oil-red-O (Millipore Sigma) staining and counter stained with Harris Hematoxylin (Millipore Sigma). Lesion size was assessed using Nanozoomer 2.0RS (Hamamatsu). Briefly, lesion area was quantified by measuring the atherosclerotic plaque of the intima from the endothelial layer to the healthy media. For GFP staining, anti-GFP antibody (1:400, ab13970, Abcam) was incubated overnight at 4°C and Alexa Fluor 488 goat anti-chicken IgY (H+L) secondary antibody (1:100, A-11039, ThermoFisher Scientific) was applied to detect GFP+ cells. The tissue sections were counterstained with DAPI (1:3000, D21490, ThermoFisher Scientific) and all the slides were scanned by a digital scanner NanoZoomer 2.0RS.

### Statistical analysis

Statistical tests and figures were generated using Prism v9.1.0. Sample sizes were chosen based on previous experiments and no statistical methods were employed to predetermine sample size. For pairwise comparisons statistical significance was calculated using unpaired, 2-tailed t-test and for comparisons between three or more groups one-way ANOVA followed by Sidak’s post-hoc analysis. All data is presented as mean ±SD and statistical significance was set to p<0.05 and significance is denoted as follows: *<0.05, **<0.01, ***<0.001 and ****<0.0001.

## Supplemental figure legends

Supplemental figure 1.

(A) Gating strategy for FACS analysis of peripheral blood subsets in mice treated with CD45-ADC and Ubiquitin-GFP HSCT. Inflammatory monocytes (Infl. monocytes); Patrolling monocytes (Patr. monocytes); Granulocytes (Granul.).

(B) Gating strategy for flow cytometric analysis of bone marrow hematopoietic stem and progenitor cells 6 weeks following CD45-ADC treatment and HSCT.

(C) Bar graphs showing absolute numbers of bone marrow hematopoietic stem and progenitors as determined by FACS analysis 6 weeks post-CD45-ADC treatment and HSCT.

Supplemental figure 2.

(A) Gating strategy for FACS analysis of brain leukocyte subsets in mice treated with CD45-ADC and Ubiquitin-GFP HSCT.

(B) Bar graphs showing absolute numbers of brain myeloid cells as determined by FACS analysis 6 weeks post-CD45-ADC treatment and HSCT.

(C) Gating strategy for FACS assessment of liver resident hematopoietic cells 6 weeks following CD45-ADC treatment and HSCT. Macrophages (Mᴓ); Kupffer cells (KC).

(D) Bar graphs summarizing the absolute numbers of liver myeloid cell types 6 weeks post-CD45-ADC treatment and HSCT.

(E) Total number of Kupffer cells and liver macrophages as determined by FACS analysis 6 weeks after CD45-ADC and HSCT summarized as bar graphs.

Supplemental figure 3.

(A) Gating strategy for flow cytometric evaluation of lung leukocytes in mice treated with CD45-ADC and Ubiquitin-GFP HSCT. MHCII High interstitial macrophages (Inter. Mᴓ^High^); (MHCII Low interstitial macrophages (Inter. Mᴓ^Low^); Alveolar macrophages (Alveolar Mᴓ); Dendritic cells (DC).

(B) Bar graphs displaying absolute numbers of lung myeloid immune cells as determined by flow cytometric analysis 6 weeks post-CD45-ADC treatment and HSCT.

(C) Gating strategy for FACS assessment of skin resident hematopoietic cells 6 weeks following CD45-ADC treatment and HSCT. Macrophages (Mᴓ); Langerhans cells (LC); Dendritic cells (DC).

(D) Bar graphs summarizing the absolute numbers of skin myeloid leukocyte types 6 weeks post-CD45-ADC treatment and HSCT.

Supplemental figure 4.

(A) Bar graphs showing absolute numbers of bone marrow hematopoietic stem and progenitors as determined by FACS analysis 6 weeks post-CD45-ADC treatment and Ubiquitin-GFP HSCT in atherosclerotic Ldlr KO mice. Megakaryocyte-erythroid progenitor (MEP); Common myeloid progenitor (CMP); Granulocyte-monocyte progenitor (GMP); Long-term hematopoietic stem cell (LT-HSC).

(B) Gating strategy for flow cytometric analysis of hematopoietic cells in aortic atherosclerotic lesions after 12 weeks of high-fat diet and 6 weeks following CD45-ADC treatment and HSCT. Macrophages (Mᴓ); Inflammatory monocytes (Infl. monocytes); Patrolling monocytes (Patr. monocytes); Granulocytes (Granul.).

(C) Bar graphs showing absolute numbers of myeloid immune cells in aortic wall atherosclerotic plaques as determined by FACS analysis 6 weeks post-CD45-ADC treatment and HSCT.

Supplemental figure 5.

(A) Validation of reduced Tet2 expression by qPCR in peripheral blood leukocytes 1 week after poly:IC induction of transplanted WT or Tet2 KO bone marrow.

(B) Bar graphs displaying CFP chimerism of LT-HSCs in atherosclerotic Ldlr KO mice transplanted with either WT or Tet2 KO bone marrow as determined by flow cytometric analysis.

(C) White blood cell counts in atherosclerotic Ldlr KO mice transplanted with either WT or Tet2 KO bone marrow, 6 weeks following CD45-ADC and HSCT.

(D) Bar graphs displaying CFP chimerism of aortic atherosclerotic lesion leukocytes in Ldlr KO mice after transplantation of either WT or Tet2 KO bone marrow as determined by flow cytometric analysis.

(E) Absolute number of cells in aortic lesions of Ldlr KO mice after transplantation of either WT or Tet2 KO bone marrow.

(F) Bar graphs showing the number of macrophages identified by flow cytometric analysis in aortic atherosclerotic lesions of Ldlr KO mice transplanted with either WT or Tet2 KO bone marrow, normalized to the lesion surface area as determined by Oil red O staining of histological sections.

## Acknowledgments

The authors thank members of D.T.S and M.N laboratories for technical assistance. We were also supported with expert technical assistance from the HSCI-CRM Flow Cytometry facility. This research was supported by the Craig A. Huff Harvard Stem Cell Institute Research Support Fund, the Gerald and Darlene Jordan Professorship of Medicine, and the NIH U19AI149676 to D.T.S. K.G was supported by the Swedish Research Council the John S. Macdougall Jr. and Olive R. Macdougall Fund.

## Conflict of Interest Statement

D.T.S. is a director and shareholder for Agios Therapeutics and Editas Medicines; a founder, director, shareholder, and scientific advisory board member for Magenta Therapeutics and LifeVault Bio, a shareholder and founder of Fate Therapeutics and Garuda Therapeutics, and a director, founder, and shareholder for Clear Creek Bio, a consultant for FOG Pharma, Inzen Therapeutics and VCanBio, and a recipient of sponsored research funding from Sumitomo Dianippon. D.T.S and K.G are inventors of patent US20220143099A1. R.P, S.L.H and A.E.B were full-time salaried employees and equity holders of Magenta Therapeutics at the time the work was completed.

