## Supplementary figures for "CD45-antibody-drug conjugate clears tissue resident myeloid cells from their niches enabling therapeutic adoptive cell transfer"

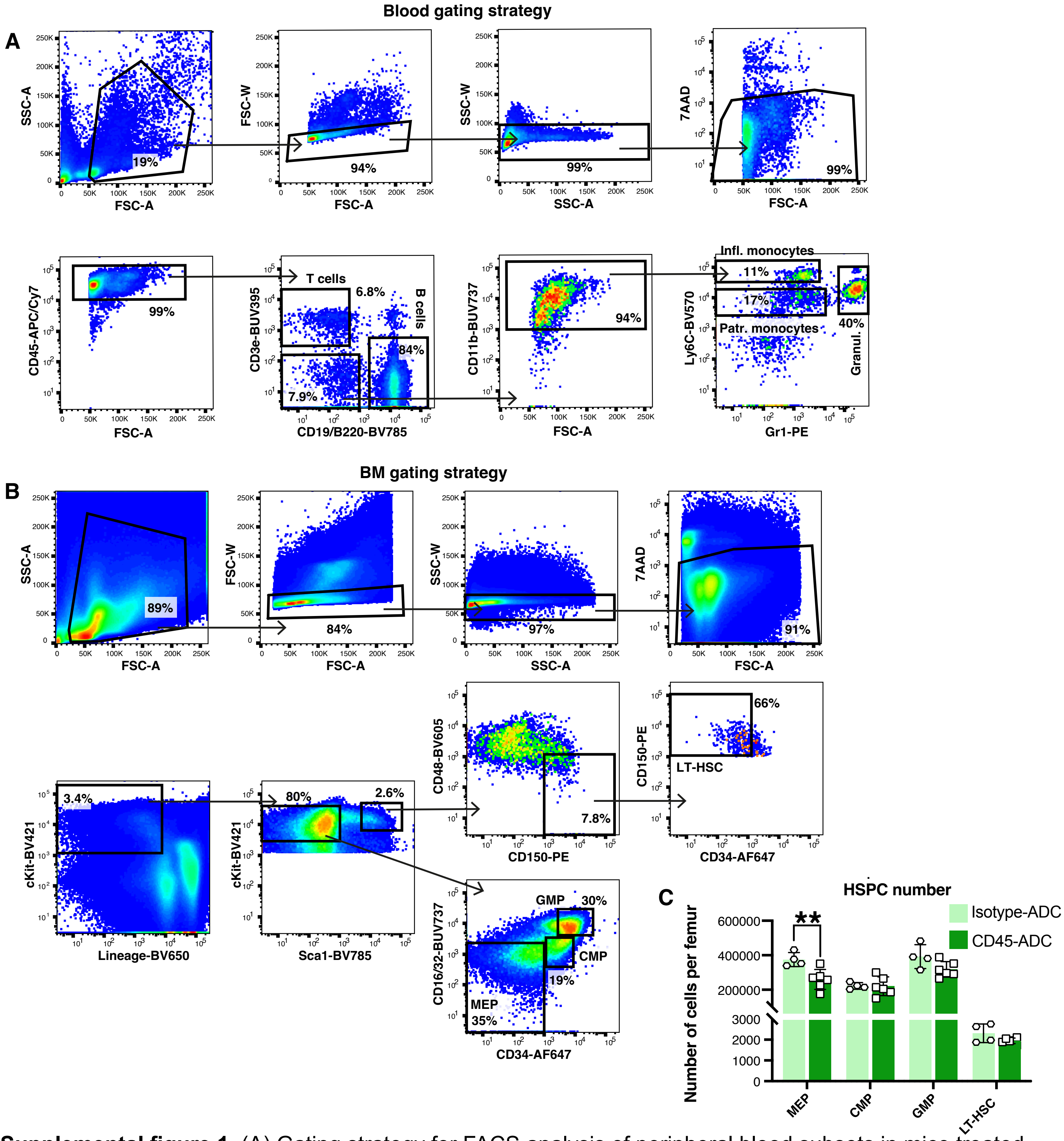


**Supplemental figure 1.** (A) Gating strategy for FACS analysis of peripheral blood subsets in mice treated with CD45-ADC and Ubiquitin-GFP HSCT. Inflammatory monocytes (Infl. monocytes); Patrolling monocytes (Patr. monocytes); Granulocytes (Granul.). (B) Gating strategy for flow cytometric analysis of bone marrow hematopoietic stem and progenitor cells 6 weeks following CD45-ADC treatment and HSCT. (C) Bar graphs showing absolute numbers of bone marrow hematopoietic stem and progenitors as determined by FACS analysis 6 weeks post-CD45-ADC treatment and HSCT.

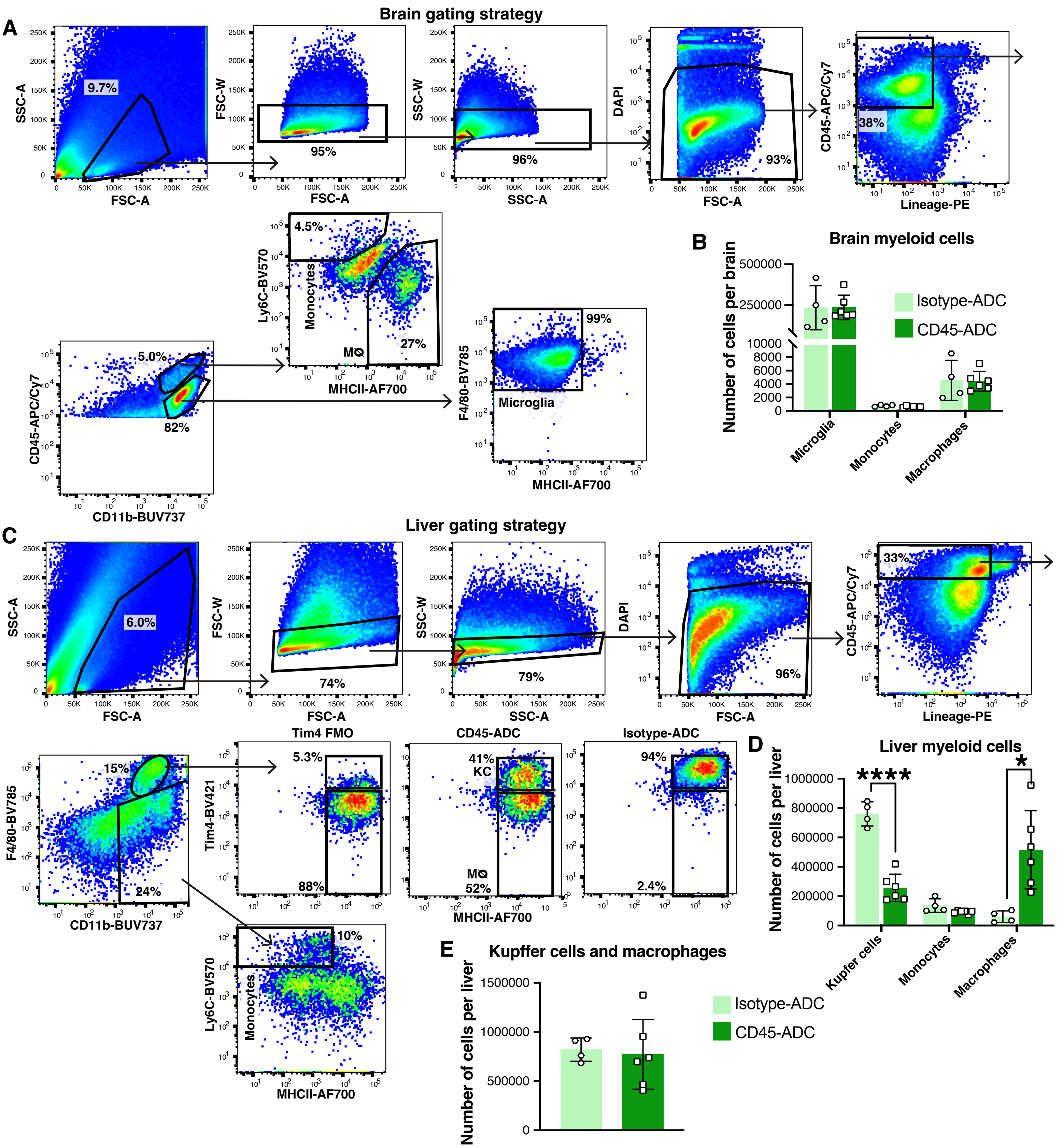


**Supplemental figure 2.** (A) Gating strategy for FACS analysis of brain leukocyte subsets in mice treated with CD45-ADC and Ubiquitin-GFP HSCT. (B) Bar graphs showing absolute numbers of brain myeloid cells as determined by FACS analysis 6 weeks post-CD45-ADC treatment and HSCT. (C) Gating strategy for FACS assessment of liver resident hematopoietic cells 6 weeks following CD45-ADC treatment and HSCT. Macrophages (Mϕ); Kupffer cells (KC). (D) Bar graphs summarizing the absolute numbers of liver myeloid cell types 6 weeks post-CD45-ADC treatment and HSCT. (E) Total number of Kupffer cells and liver macrophages as determined by FACS analysis 6 weeks after CD45-ADC and HSCT summarized as bar graphs.

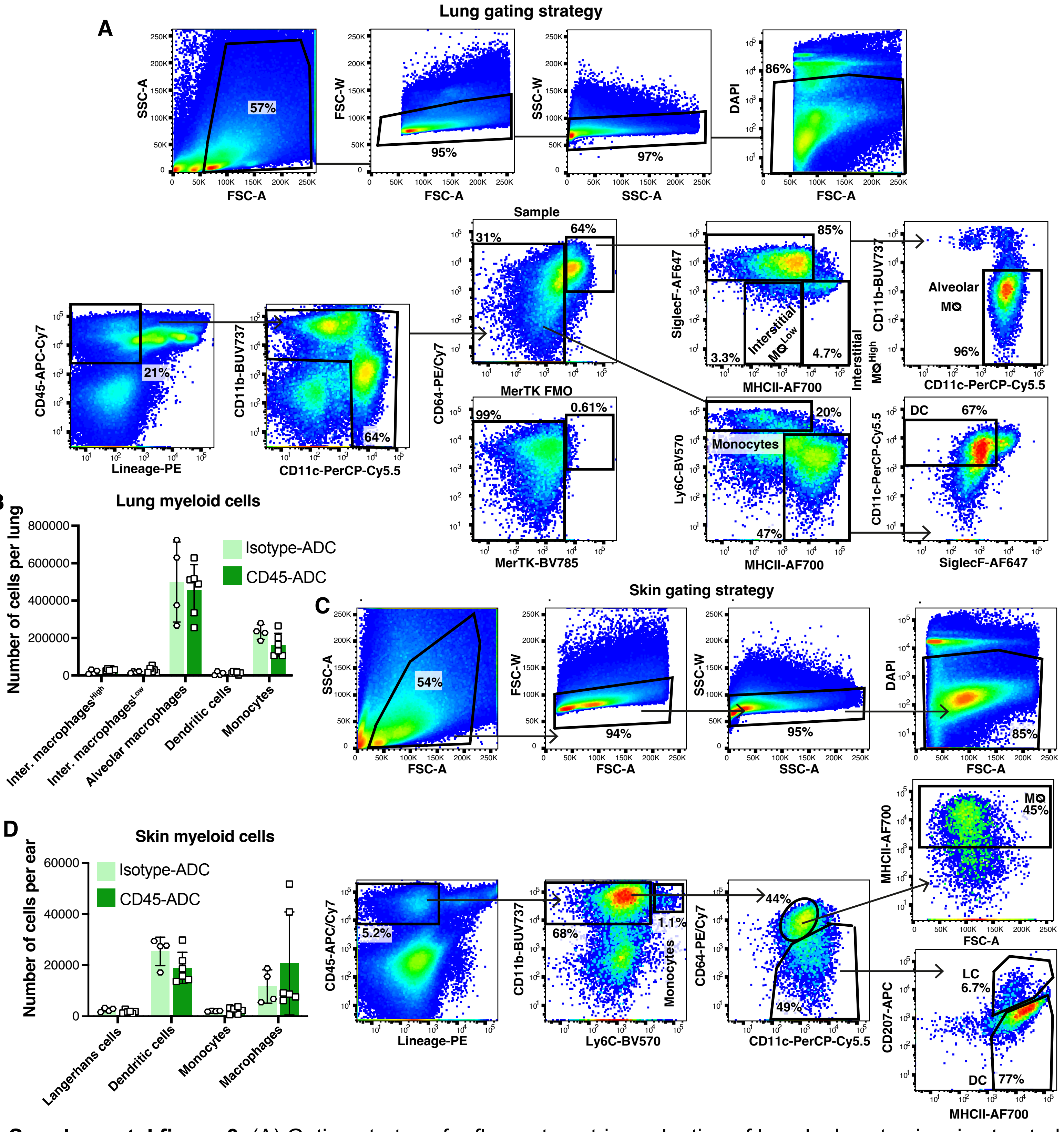


**Supplemental figure 3.** (A) Gating strategy for flow cytometric evaluation of lung leukocytes in mice treated with CD45-ADC and Ubiquitin-GFP HSCT. MHCII High interstitial macrophages (Inter. MΦHigh); (MHCII Low interstitial macrophages (Inter. MΦLow); Alveolar macrophages (Alveolar MΦ); Dendritic cells (DC). (B) Bar graphs displaying absolute numbers of lung myeloid immune cells as determined by flow cytometric analysis 6 weeks post-CD45-ADC treatment and HSCT. (C) Gating strategy for FACS assessment of skin resident hematopoietic cells 6 weeks following CD45-ADC treatment and HSCT. Macrophages (MΦ); Langerhans cells (LC); Dendritic cells (DC). (D) Bar graphs summarizing the absolute numbers of skin myeloid leukocyte types 6 weeks post-CD45-ADC treatment and HSCT.

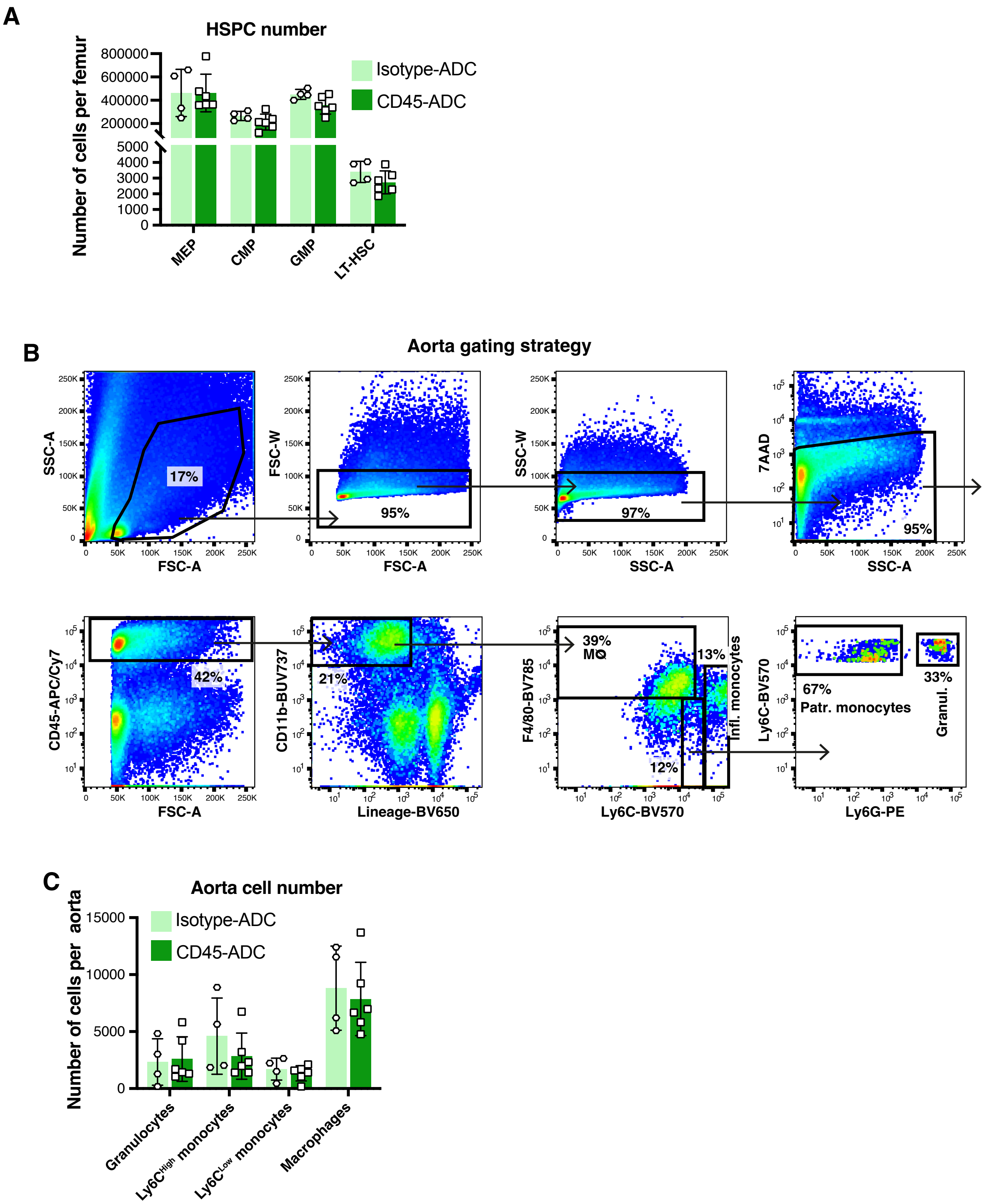


**Supplemental figure 4.** (A) Bar graphs showing absolute numbers of bone marrow hematopoietic stem and progenitors as determined by FACS analysis 6 weeks post-CD45-ADC treatment and Ubiquitin-GFP HSCT in atherosclerotic Ldlr KO mice. Megakaryocyte-erythroid progenitor (MEP); Common myeloid progenitor (CMP); Granulocyte-monocyte progenitor (GMP); Long-term hematopoietic stem cell (LT-HSC). (B) Gating strategy for flow cytometric analysis of hematopoietic cells in aortic atherosclerotic lesions after 12 weeks of high-fat diet and 6 weeks following CD45-ADC treatment and HSCT. Macrophages (MΦ); Inflammatory monocytes (Infl. monocytes); Patrolling monocytes (Patr. monocytes); Granulocytes (Granul.). (C) Bar graphs showing absolute numbers of myeloid immune cells in aortic wall atherosclerotic plaques as determined by FACS analysis 6 weeks post-CD45-ADC treatment and HSCT.

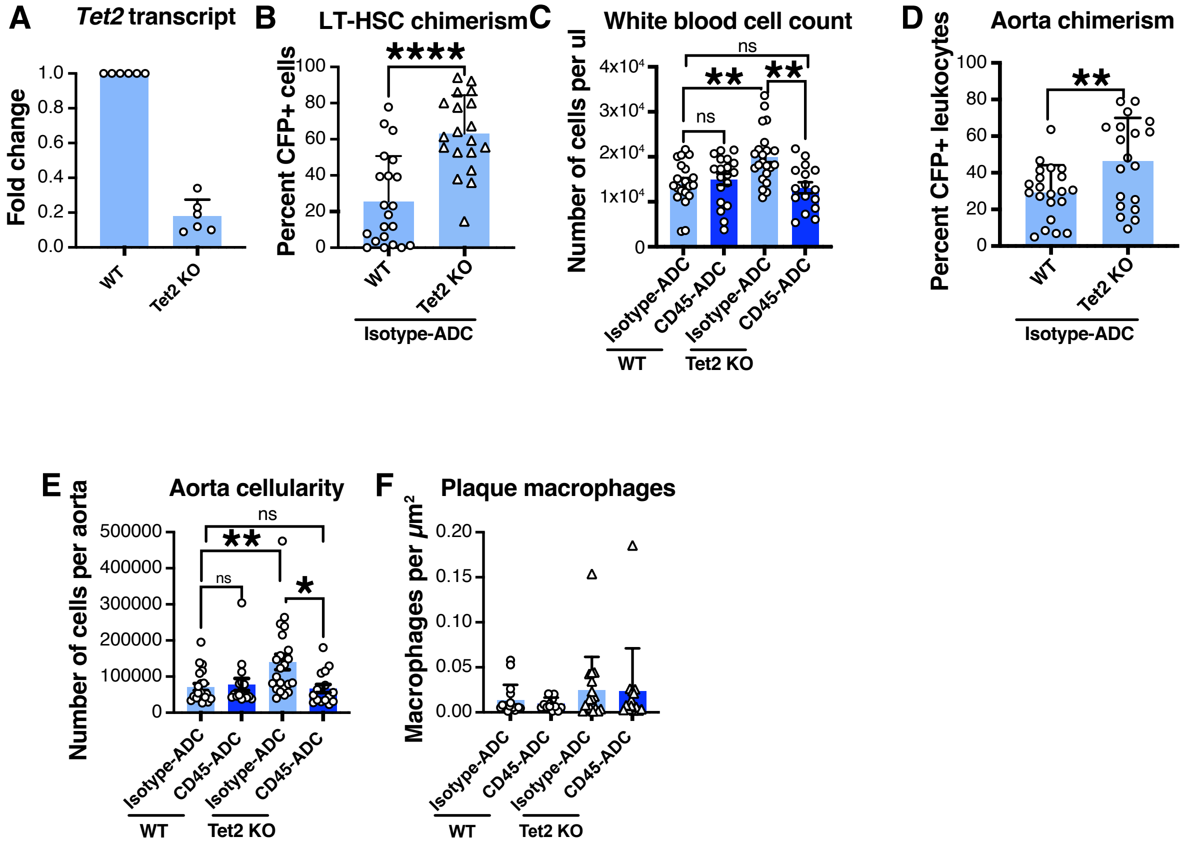


**Supplemental figure 5.** (A) Validation of reduced Tet2 expression by qPCR in peripheral blood leukocytes 1 week after poly:IC induction of transplanted WT or Tet2 KO bone marrow. (B) Bar graphs displaying CFP chimerism of LT-HSCs in atherosclerotic Ldlr KO mice transplanted with either WT or Tet2 KO bone marrow as determined by flow cytometric analysis. (C) White blood cell counts in atherosclerotic Ldlr KO mice transplanted with either WT or Tet2 KO bone marrow, 6 weeks following CD45-ADC and HSCT. (D) Bar graphs displaying CFP chimerism of aortic atherosclerotic lesion leukocytes in Ldlr KO mice after transplantation of either WT or Tet2 KO bone marrow as determined by flow cytometric analysis. (E) Absolute number of cells in aortic lesions of Ldlr KO mice after transplantation of either WT or Tet2 KO bone marrow. (F) Bar graphs showing the number of macrophages identified by flow cytometric analysis in aortic atherosclerotic lesions of Ldlr KO mice transplanted with either WT or Tet2 KO bone marrow, normalized to the lesion surface area as determined by Oil red O staining of histological sections.
